# Thermal sensitivity of the *Spiroplasma-Drosophila hydei* protective symbiosis: The best of climes, the worst of climes

**DOI:** 10.1101/2020.04.30.070938

**Authors:** Chris Corbin, Jordan E. Jones, Ewa Chrostek, Andy Fenton, Gregory D. D. Hurst

## Abstract

The outcome of natural enemy attack in insects has commonly been found to be influenced by the presence of protective symbionts in the host. The degree to which protection functions in natural populations, however, will depend on the robustness of the phenotype to variation in the abiotic environment. We studied the impact of a key environmental parameter – temperature – on the efficacy of the protective effect of the symbiont *Spiroplasma* on its host *Drosophila hydei*, against attack by the parasitoid wasp *Leptopilina heterotoma*. In addition, we investigated the thermal sensitivity of the symbiont’s vertical transmission, which may be a key determinant of the ability of the symbiont to persist. We found that vertical transmission was more robust than previously considered, with *Spiroplasma* being maintained at 25 °C, 18 °C and with 18/15 °C diurnal cycles, with rates of segregational loss only increasing at 15 °C. Protection against wasp attack was ablated before symbiont transmission was lost, with the symbiont failing to rescue the fly host at 18 °C. We conclude that the presence of a protective symbiosis in natural populations cannot be simply inferred from presence of a symbiont whose protective capacity has been tested under narrow controlled conditions. More broadly, we argue that the thermal environment is likely to represent an important determinant of the evolutionary ecology of defensive symbioses in natural environments, potentially driving seasonal, latitudinal and altitudinal variation in symbiont frequency, and modulating the strength of selection for symbiotic protective systems compared to defensive systems encoded in the nuclear genomes.

## Introduction

It is now recognised that resistance to natural enemy attack in insects is commonly determined by protective symbionts (Oliver *et al.*, 2014). Maternally-transmitted bacterial symbionts defend their hosts against attacks, which represents an adaptation by the symbiont to promote its own vertical transmission. Defence occurs through either competition with the natural enemy, the direct production of toxins that kill or inhibit the parasite, or indirectly through modulating the host immune system (Gerardo and Parker, 2014).

These symbioses are most well understood in aphid systems, where a variety of heritable bacteria defend their hosts against attack by fungi and wasps (Oliver *et al.*, 2003; Scarborough *et al.*, 2005). However, protective symbiosis is more widely observed across insects. In *Drosophila*, for instance, resistance to viral attack is commonly determined by the presence of *Wolbachia* (Hedges *et al.*, 2008; Teixeira *et al.*, 2008) and the ability to survive attack by wasps and nematodes is a function of the presence of *Spiroplasma* (Xie *et al.*, 2010; Jaenike *et al.*, 2010). That heritable microbes represent part of the adaptive response of a host population to attack is evidenced by rapid spread of the defensive symbiont in response to parasite pressure both in laboratory culture and in the field (Oliver *et al.*, 2008; Jaenike *et al.*, 2010; Jaenike and Brekke, 2011; Xie *et al.*, 2015).

Our understanding of the evolutionary ecology of defensive symbiosis has emphasised genetic properties of the host, symbiont and natural enemy that determine the outcome of a parasite-host encounter (Parker *et al.*, 2017), with strong interaction terms leading to predictions of complex cyclical dynamics (Vorburger and Perlman, 2018). There is also an awareness that other aspects of the biotic environment may also be dynamically important. In the aphid system, for example, the host plant upon which an aphid feeds, and the presence of other plant species, may alter the protective phenotype (Sochard *et al.*, 2019).

There has however been less study of the impact of the abiotic environment on protective symbioses. There are two reasons to believe that thermal environment, in particular, will alter both individual and population level outcomes of protective symbiosis. First, the outcome of parasite-host interactions in poikilotherms is commonly sensitive to temperature (Thomas and Blanford, 2003). Endogenous host defences (that may combine with symbiont-mediated protection) will commonly vary with temperature, as will parasite infectivity and virulence traits, and the interaction of these may change the outcome of attack. Second, other symbiont-mediated traits are commonly thermally sensitive (Corbin *et al.*, 2017). Thermal impacts on either the strength of protection at the individual level, or vertical transmission efficiency, would alter both the frequency of protected individuals in a population and the degree of protection afforded when they are present. Studies of thermal impact on natural protective symbiotic associations to date are limited to aphid systems, where either high temperatures or transient heat shock reduce protection (Heyworth and Ferrari, 2016; Doremus *et al.*, 2018). Their broader effects for other insect host – symbiont – enemy interactions are currently not known.

In this study, we examine thermal sensitivity in the *Drosophila hydei*-*Spiroplasma* protective symbiosis. *Spiroplasma* heritable symbionts infect a wide variety of *Drosophila* species (Watts *et al.*, 2009), and display two main phenotypes, male-killing and protection, that may exist alone or in combination (Montenegro *et al.*, 2005; Xie *et al.*, 2010; Jaenike *et al.*, 2010). Past surveys have shown 23-66% of *D. hydei* carry *Spiroplasma* (Ota *et al.*, 1979; Kageyama *et al.*, 2006; Watts *et al.*, 2009; Osaka *et al.*, 2013). Two *Spiroplasma* strains exist in *D. hydei* (Mateos *et al.*, 2006), of which one (Hy1, also referred to as sHy1) is most common. Experiments have demonstrated that the Hy1 strain confers protection against attack by *Leptopilina heterotoma* parasitoid wasps (Xie *et al.*, 2010). Studies in Japanese populations of this fly indicated that vertical transmission was temperature sensitive, with no transmission at 15 °C, substantial segregational loss at 18 °C and high vertical transmission at 25 °C (Osaka *et al.*, 2008). *Spiroplasma* titre was greatly reduced when flies were reared at 18 °C compared to 25 °C. These data, coupled to evidence of low temperature cure and loss of male-killing phenotype for *Spiroplasma poulsonii* in *D. melanogaster* (Montenegro and Klaczko, 2004), indicate the thermal sensitivity of these symbioses, in particular the potential for low temperature to ablate the phenotype and ultimately cure infection.

The current data leave two questions. First, how are *Spiroplasma* maintained in temperate populations of *D. hydei* if 15 °C produces an absolute cure? Second, are protective phenotypes thermally sensitive, like male-killing phenotypes? Our approach to assess the impact of thermal environment on the protective symbiosis is threefold. Previous survey work examining *Spiroplasma* presence has been conducted in warm temperate regions (below 36° N latitude) (Mateos *et al.*, 2006; Watts *et al.*, 2009; Osaka *et al.*, 2013). To investigate if the symbiosis was present at cooler latitudes, we complemented this with estimation of *Spiroplasma* prevalence in *D. hydei* from the southern UK (51° N), which is towards the northern end of the species range. Following this, we examined the impact of temperature on *Spiroplasma* vertical transmission over two vertical transmission events. Finally, we investigate the impact of thermal environment on the defence phenotype, protection against attack by *L. heterotoma*. Our experimental conditions here mimic cool/warm environments, with a generation maintained at 18°C or 25°C producing progeny that are then also challenged at the focal temperature. These results show that whilst transmission of the focal strain of *Spiroplasma* is more robust to thermal environment than previously reported, but the protection phenotype is ablated at low temperatures.

## Methods

### Prevalence estimates for *Spiroplasma* in UK *D. hydei*

Wild *D. hydei* specimens were collected in Royal Tunbridge Wells (51.09 N, 0.16 E), in July 2014 and August 2015. Adult flies were caught with sweep nets over fruit bait and transferred to vials containing SY food (Sugar Yeast Medium: 20 g agarose, 100 g sugar, 100 g autolysed yeast in a total volume of 1 L, to which 30 mL 10% Nipagin w/v propionic acid was added). The nearest climate station to the collecting site is ‘Herstmonceux West End’, where the average minimum temperature during August over the years 1981-2010 was 12.7 °C and the average maximum temperature was 21.5 °C (Met Office, https://www.metoffice.gov.uk/research/climate/maps-and-data/uk-climate-averages/u101x20r9).

Collected flies were kept live in a CT room at 25 °C for 2 weeks to reduce false negative PCR assays arising from low titre, before being frozen at -80 °C. DNA was prepared through the Promega Wizard® protocol, and the template quality was tested using COI amplification. PCR amplifications were carried out using GoTaq Hot Start Green Master Mix (Promega). Templates passing quality control were tested for the presence of *Spiroplasma* by PCR assay, using the primer combination SpoulF and SpoulR (henceforth, *Spiroplasma* PCR assay) (Montenegro *et al.*, 2005) (see SOM Table S1).

### Strains used in experiments on protection and transmission

Protection, titre and transmission assays utilized a strain derived from Mexican *D. hydei* stock (TEN104-106) originally established from a single *Spiroplasma* strain sHy1-infected female (Mateos *et al.*, 2006) and maintained since this time at 25 °C. An uninfected stock was subsequently produced through tetracycline-treatment. Shortly before this experiment (in May 2014), a new isogenic infected line was generated by injecting haemolymph from adult female donors of the original infected line, into adult female flies from the cured line. The injected flies were bred on ASG vials (Corn Meal Agar: 10 g agarose, 85 g sugar, 60 g maize meal, 40 g autolysed yeast in a total volume of 1 L, to which 25 mL 10% Nipagin was added). Offspring from females positive in the *Spiroplasma* PCR assay were used to re-establish the infected line.

Assays on transmission additionally utilized a strain collected in Cambridge, UK. The Cambridge strain was established from a single, sHy1 positive mated female caught in Cambridge in September 2013, and maintained on ASG vials at 25 °C until early 2014, when it was transferred to cornmeal agar bottles to increase numbers. The identity of *Spiroplasma* Hy1 was confirmed through sequence of partial 16S rRNA gene sequence.

Protection assays utilised an inbred *Leptopilina heterotoma* wasp strain collected from Sainte Foy-lès-Lyon and la Voulte, France (Jones and Hurst, 2020).

### Thermal effects on transmission

We tested the vertical transmission efficiency of two host-symbiont combinations through passage across a range of temperatures, maintaining a population of flies, and determining *Spiroplasma* presence in third generation (F3) adults, following two vertical transmission events, within these populations.

To homogenise rearing conditions of the Cambridge and Mexican flies prior to the experiment, at 25 °C, the mothers of generation P laid eggs in large ASG vials containing an adult male. After two days, the flies were tipped into a new vial for an additional two days, before females were individually frozen at -80 °C. The mothers underwent DNA extraction (Promega Wizard® kit) and were tested for *Spiroplasma* using hot-start PCR with SpoulF/SpoulR primers. Only vials from sHy1 infected mothers were kept and matured. The progeny of these infected mothers, generation P, were matured for 13-17 days at 25 °C.

For each line, two population cages were established at 25 °C to generate F1 larvae. Each cage contained 50 female and 10 male generation-P flies on a grape-juice agar plate ‘painted’ with live yeast paste, which was replaced daily. Once larvae were successfully obtained on plates, 20 breeding females from each cage were frozen at -80 °C for later verification of infection status. After ageing for a day, larvae were picked from plates into 75 mm vials containing 8 ml of ASG food at a density of 25 larvae per vial. Sixteen vials were picked from each plate, giving 32 vials per line. Vials were then randomly-assigned to one of four temperature conditions (Constant 25 °C, 18 °C, 15 °C, or day night alternating 18/15 °C), using the shuffle function in statistical software R (R Core Team, 2013), such that 8 vials were placed in each temperature condition for each line. Larval vials were watered and shuffled in their storage tray twice a week.

The lines were reared to eclosion, which took approximately 2 weeks at 25 °C, 3 weeks at 18 °C, and 4-5 weeks at 18/15 °C and 15 °C. Flies were sexed within 3-4 days of eclosion and maintained on SY food. After 3, 7, 7 and 9 days at temperatures 25 °C, 18 °C, 18/15 °C and 15 °C respectively, the experimental-temperature adult females from each vial of origin were mated to stock *Spiroplasma*-negative males and left to lay eggs in individual vials at their experimental temperature. The laying females were tipped onto a new vial so that they produced two vials of eggs in total. Vial 1 was kept at experimental temperature, while vial 2 was transferred to 25 °C, to increase titre of any transmitted *Spiroplasma* for detection. The female was then frozen at -80 °C for verification of infection status.

Once eclosed and mature, F2s at experimental temperatures were bred from, while F2s at the control temperature were frozen on maturity without breeding. F3 eggs obtained from F2 experimental temperature crosses were picked into vials and exposed to their respective thermal environments. Upon eclosing and being permitted to mature, they were allowed to recover as adult at 25 °C to increase titre, and frozen for PCR testing without being bred from.

### DNA extraction and PCR assay for *Spiroplasma*

For experimental flies, DNA was extracted by homogenising whole flies in a mix of 5% w/v Chelex 100 in molecular-grade water (BioRad) with 2 µl proteinase K (20 mg.ml), incubating at 37 °C overnight, then centrifuging to produce DNA-containing supernatant. DNA samples which tested negative for *Spiroplasma* were subsequently ‘cleaned up’ using a modified protocol for the Wizard® DNA Extraction Kit (Promega), to reduce inhibition of PCR by substances in the fly, before undergoing PCR test again.

### Thermal influences on protection

Our approach was to emulate protection during protracted cool vs warm periods, reflecting changes in *Spiroplasma* titre associated with longterm exposure to the focal temperature, attack by wasps occurring at that temperature (with the period of exposure to parasites mirroring the length of the susceptible L2/L3 stage) and defence also occurring at that temperature.

Resistance of flies to wasp attack, and the capacity of wasps to complete development, were compared at 18 °C and 25 °C, for *Spiroplasma* infected and uninfected flies, following a generation of passage at the focal temperature. A 2×2×2 factorial design was completed, examining fly survival in the presence or absence of wasps, presence or absence of *Spiroplasma*, and at 18 °C or 25 °C. Constraints on the numbers of fertile female *L. heterotoma* available at any one time led to the experiment being split into two blocks, Block A and Block B, repeated 2-3 months apart in the same incubators and under the same conditions. The block design was incorporated into later statistical analyses. Attack of flies occurs during the second larval instar, and protection was tested in the progeny of flies reared at the focal temperature.

sHy1-positive stock and sHy1-negative grandparent stocks of the TEN-104 line previously maintained at 25 °C laid eggs in separate bottles at both 25 °C and 18°C (4 bottles total, one of each temperature/hy1 status combination). Each bottle contained ∼50 females and ∼20 males. Flies laid eggs for 2 days at 25 °C and 4 days at 18 °C, then adults were disposed of to prevent generations mixing. Parent stocks were reared in 12 hour/12 hour light/dark cycle incubators at the focal temperature (Sanyo MLR-351), sexed on their eclosion days, and the females stored at their birth temperatures on SY food.

After reaching sexual maturity (day 2 at 25 °C and day 4 at 18 °C) the female flies created above were placed in individual population cages over a small yeast-painted grape juice agar plate, each with two sHy1-negative Mexican isoline males which were at least 6 days old. Females were permitted to mate and lay eggs for one day at 25 °C and two days at 18 °C, and tipped onto new plates after this period, repeating until sufficient larvae were obtained. Three days after laying commenced at 25 °C, and 6 days after laying commenced at 18 °C, L1 larvae were picked with hooks onto small ASG vials, such that the larvae in a vial all came from one known mother. Target larval density was 15 larvae per vial, but due to laying rate constraints, some vials contained 7 or 8 larvae. Mothers of vials were frozen at -80 °C for later infection status verification by PCR assay. Only larvae from verified-infected mothers were included in the experiment.

After picking, larvae were immediately exposed to *L. heterotoma* wasps. The wasps had previously been matured to at least 7 days of age at 22 °C on grape agar vials, with honey available for nutrition, and then given three days of oviposition experience on L1-L2 *Drosophila melanogaster* (Oregon-R). Five female wasps and three males were transferred to each picked vial of larvae; pilot attempts (data not shown) had used three female wasps and three male wasps per vial, but this had been insufficient for a good attack rate. The wasps were left to attack larvae over the L2/L3 larval period (three days at 25 °C and six days at 18 °C), emulating the period of fly development at which wasps parasitize in the field. This protocol led to an attack rate of >80% at both temperatures (fewer than 20% of flies emerge in the parasitisation/no symbiont treatment vs >95% emergence in absence of parasite).

Vials were then monitored daily. The numbers of eclosing flies and wasps were counted for each vial. Typically, at 25 °C, fly emergence began at day 14 and wasp emergence at day 21. These times were approximately doubled at 18 °C. Observations continued until 30 days after picking at 25 °C, and 60 days after picking at 18 °C.

The impact of this rearing regime on *Spiroplasma* titre was also assessed. Flies carrying *Spiroplasma* were again reared through the focal temperature, and F1 progeny obtained. These flies were then culled on entry to pupation, as this represents the time of active engagement between wasp and *Spiroplasma.* DNA template was prepared from individual pupae, and *Spiroplasma* titre estimated by qPCR. To this end, total DNA was extracted from 10-15 individual *D. hydei* pupae per condition per replicate using the phenol-chloroform method. DNA concentrations and quality were measured with NanoDrop ND-1000 Spectrophotometer. Real-time qPCR reactions (see SOM Table 1) were then carried out for the *dnaA* gene (*Spiroplasma*) and the *rp49* gene (*D. hydei* reference), using the LightCycler 480 (Roche) (Anbutsu and Fukatsu, 2003; Granzotto *et al.*, 2011). Each reaction consisted of 6 µl of *Power* SYBR(tm) Green PCR Master Mix (ThermoFisher), 0,5 µl of each primer solution at 3.6 µM and 5 µl of diluted DNA, with three technical replicates per reaction. Melting curves were analysed to confirm specificity of amplified products. Samples scored as uninfected based on melting curve analysis were removed. Relative amounts of *Spiroplasma* were calculated using the Pfaffl Method (Pfaffl, 2001).

**Table 1:** *Spiroplasma* prevalence data from Tunbridge Wells, U.K., by year and fly sex.

|  | 2014 |  | 2015 |  |
| --- | --- | --- | --- | --- |
|  | **Males** | **Females** | **Males** | **Females** |
| Infected | **6** | **3** | **15** | **12** |
| Uninfected | **24** | **27** | **78** | **78** |

### Analysis

All statistics were carried out in R version 3.0.2 (R Core Team, 2013). The measure of fly fitness was the number of flies eclosing, divided by the number from which wasps emerged, plus pupal deaths. Wasp fitness was the number of pupae from which wasps eclosed, versus number of puparia from which flies emerged, plus pupal deaths. The non-wasp-attacked control group data was included in statistical analysis for the flies, but not for the wasps, for whom it would have been uninformative. The glm() function was used to carry out a binomial GLM on fly fitness and wasp fitness. The maximal model contained temperature, symbiont infection, wasp attack (fly fitness data only), the interactions between these three factors, plus block. The functions drop1() and update() were used to refine the model.

Relative *Spiroplasma* abundance qPCR data were tested for normality with Shapiro-Wilk test, and subsequently analysed with Kruskal-Wallis rank sum test.

## Results

### Summertime prevalence of *Spiroplasma* in *D. hydei* in the south of England

Nine out of 60 wild-caught flies (15%) from Tunbridge Wells in July 2014, and 27 of 183 flies (14.75%) collected in August 2015, tested positive for *Spiroplasma* (Table 1). Partial 16S rRNA sequence was obtained for 25 *Spiroplasma* positive flies and these were all confirmed as carrying the sHy1 strain of *Spiroplasma*. There was no evidence of heterogeneity in the proportions infected by year of collection or host sex (binomial GLM p>0.05 for year, sex and year:sex interaction). Data were therefore combined, producing a prevalence estimate of 14.8 % flies carrying *Spiroplasma* (binomial CI 10.6%-19.9%).

### Thermal impacts on symbiont vertical transmission

The symbiont persisted to the F3 generation for both *Spiroplasma*/fly isoline combinations tested (Mexico and Cambridge, henceforth ‘strains’) under all thermal environments: constant 25 °C, 18 °C, 15 °C, or day/night alternating 18/15 °C (Figure 1). A binomial GLM was used to compare infection frequency amongst F3 generation flies from the different temperature groups that had experienced two generations of transmission at experimental temperatures before being returned to 25 °C to enable proliferation of *Spiroplasma* before assay. In the final model, temperature (P < 0.001), strain (Cambridge vs Mexico) (P = 0.0074) and the temperature-strain interaction (P < 0.001) were significant factors (Table S2).

**Figure 1:**
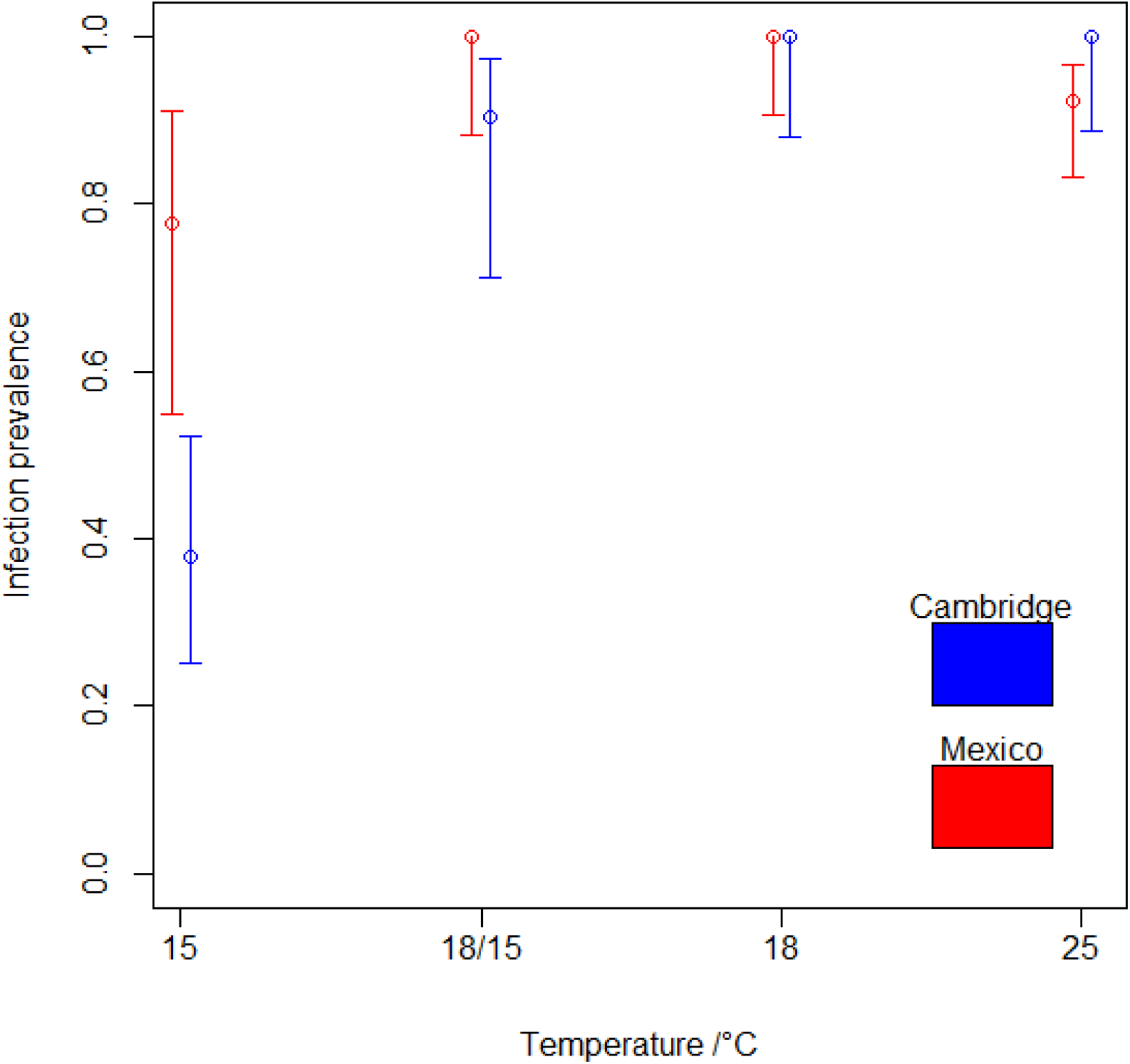
Prevalence of *Spiroplasma* infection following three generations of passage across four different thermal environments, given for each of the two different isolines. “18/15” represents alternating day/night environments respectively. Error bars represent 95% confidence intervals, calculated using the Wilson method. Sample sizes: Cambridge: 15°C N=45; 18/15°C N=19; 18 °C N=28; 25 °C N= 30; Mexico: 15°C N=18; 18/15°C N=29; 18 °C N=37; 25 °C N= 65.

Considering both the Mexico and Cambridge strains individually (see Figure 1), the confidence intervals for the 25 °C, 18 °C and 18/15 °C groups all overlap each other’s prevalence values, while this is not the case for the prevalence at 15 °C. Thus, the significance of temperature is driven by prevalence differences between the 15 °C condition and the other thermal environments.

When comparing the strains, prevalence was similarly high at 25 °C, 18 °C and 18/15 °C. However, *Spiroplasma* frequency in the Cambridge strain was significantly lower at the F3 following passage at 15 °C compared to the Mexico strain, causing the temperature-strain interaction in the full model.

### Thermal impacts on *Spiroplasma*-mediated resistance to wasp attack

The number of emerging flies was scored in the presence and absence of attacking wasps, and the presence and absence of *Spiroplasma*, for larvae of flies whose mothers had been reared 18 °C and 25 °C (Figure 2). Wasp attack rates were high at both temperatures, with emergence rates of flies below 20% in the absence of symbionts. For fly fitness, the minimal model was Fitness ∼ Temp + Inf. + Attack + Temp:Inf (Table S3). The probability of fly emergence was significantly affected by symbiont infection (p = 0.0062), the temperature x symbiont infection interaction (p = 0.0011) and the presence of wasps (p < 0.001). Temperature alone was not significant but was retained in the model because of its interaction with symbiont infection. Block was removed during the model-refining process as this didn’t significantly increase the model AIC.

**Figure 2:**
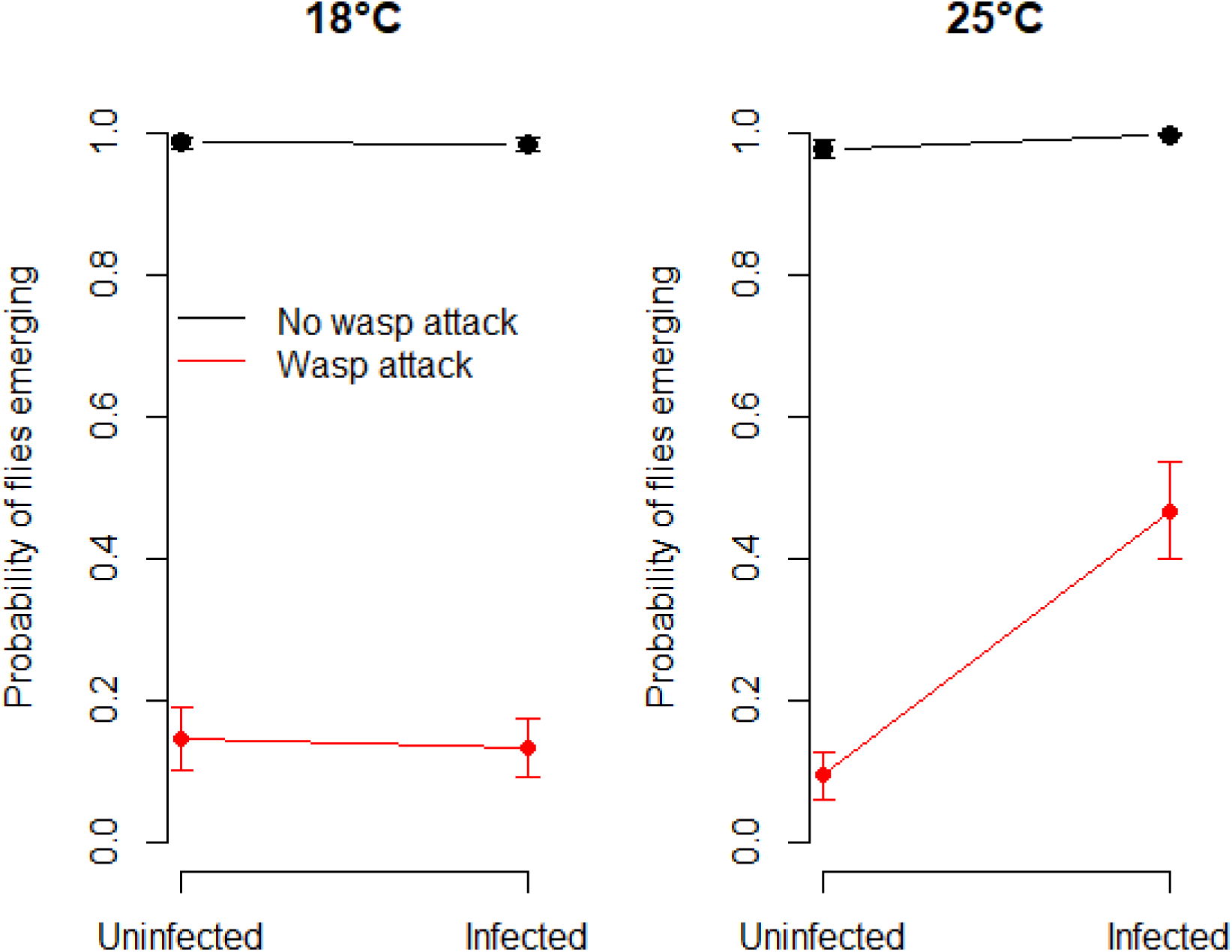
Probability of fly emergence under varying conditions of wasp attack, *Spiroplasma* infection status, and temperature. Values are as predicted from the final GLM. Error bars show the standard error of the predicted values. Sample sizes, in the format ‘total number of pupae (total number of replicate vials)’: **18°C S- Lh-**, 83 (8); **18°C S+ Lh-**, 95 (8); **18°C S- Lh+**, 56 (8); **18°C S+ Lh+**, 57 (8); **25°C S- Lh-**, 84 (7); **25°C S+ Lh-**, 94 (8); **25°C S- Lh+**, 64 (9); **25°C S+ Lh+**, 54 (9).

The significance of the symbiont infection × temperature interaction term reflects the survival of *Spiroplasma*-infected flies following parasitic wasp attack at 25 °C, compared to the absence of any discernible effect of *Spiroplasma* on fly survival following wasp attack at 18 °C (Figure 2). At 18 °C, survival in symbiont infected flies attacked by wasps was no different from uninfected flies, indicating that sHy1’s protective phenotype is depressed at the cooler temperature.

Whilst *Spiroplasma* did not aid fly survival at 18 °C, it did nevertheless impact the ability of the wasp to complete its life cycle, as measured by the number of emerging wasps (Figure 3). For wasp fitness, the minimal model was Fitness ∼ Temp + Inf. + Block (Table S4). The probability of wasp emergence was significantly affected by symbiont infection (p < 0.001), due sHy1’s protective phenotype, and temperature (p < 0.001) as the probability of wasp emergence was higher at 18 °C compared to 25 °C. Wasp emergence was generally inhibited by *Spiroplasma* presence. However, specific influence of *Spiroplasma* on wasp emergence according to temperature was not detected (a Temperature x Symbiont Infection interaction term was dropped to increase model stability), in contrast to analyses for fly survival. Experimental block was also retained in the minimal model for wasp emergence (p = 0.025), with wasp emergence generally being lower in block B (Figure 3).

**Figure 3:**
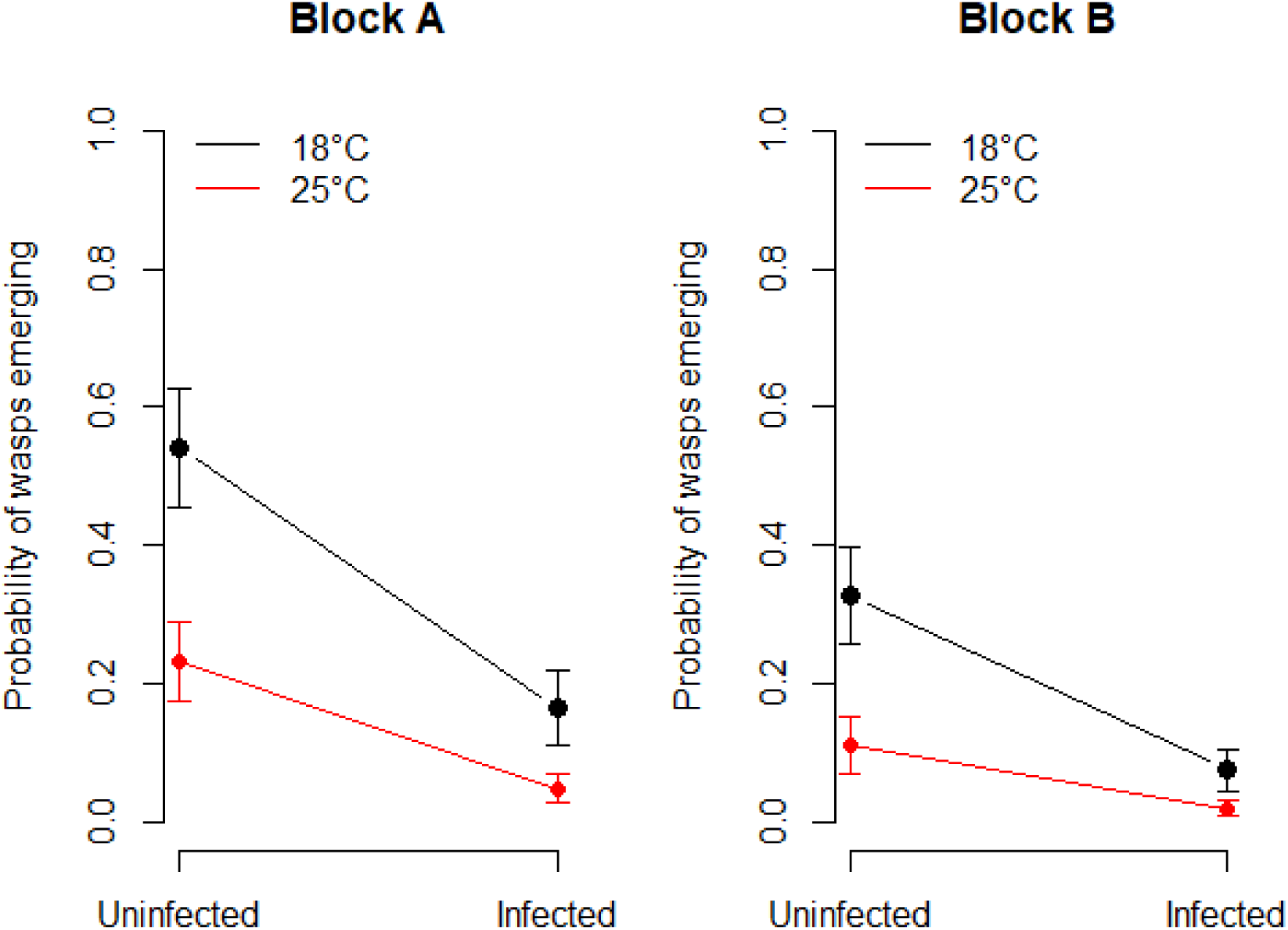
Probability of wasp emergence in wasp-attacked vials by infection status, temperature, and block. Values predicted from the final GLM. Error bars show the standard error of the predicted values. Sample sizes, in the format ‘total number of pupae (total number of replicate vials)’ for Block A: **18°C S-**, 22 (3); **18°C S+**, 29 (3); **25°C S-**, 41 (5); **25°C S+**, 36 (5). Sample sizes for Block B: **18°C S-**, 34 (5); **18°C S+**, 28 (5); **25°C S-**, 21 (4); **25°C S+**, 18 (4).

qPCR established that the thermal environment conditions imposed did impact *Spiroplasma* titre in the expected fashion (Figure 4). Pupae from the 25 °C treatment had over 3.6 times higher *Spiroplasma* titre compared to those from 18 °C treatment (Block A: 3.6x higher, Kruskal Wallis test: χ^2^ = 4.73 1 d.f. p = 0.03; Block B: 11.9x higher, Kruskal Wallis test χ^2^ = 4.77, 1 d.f. p = 0.03).

**Figure 4:**
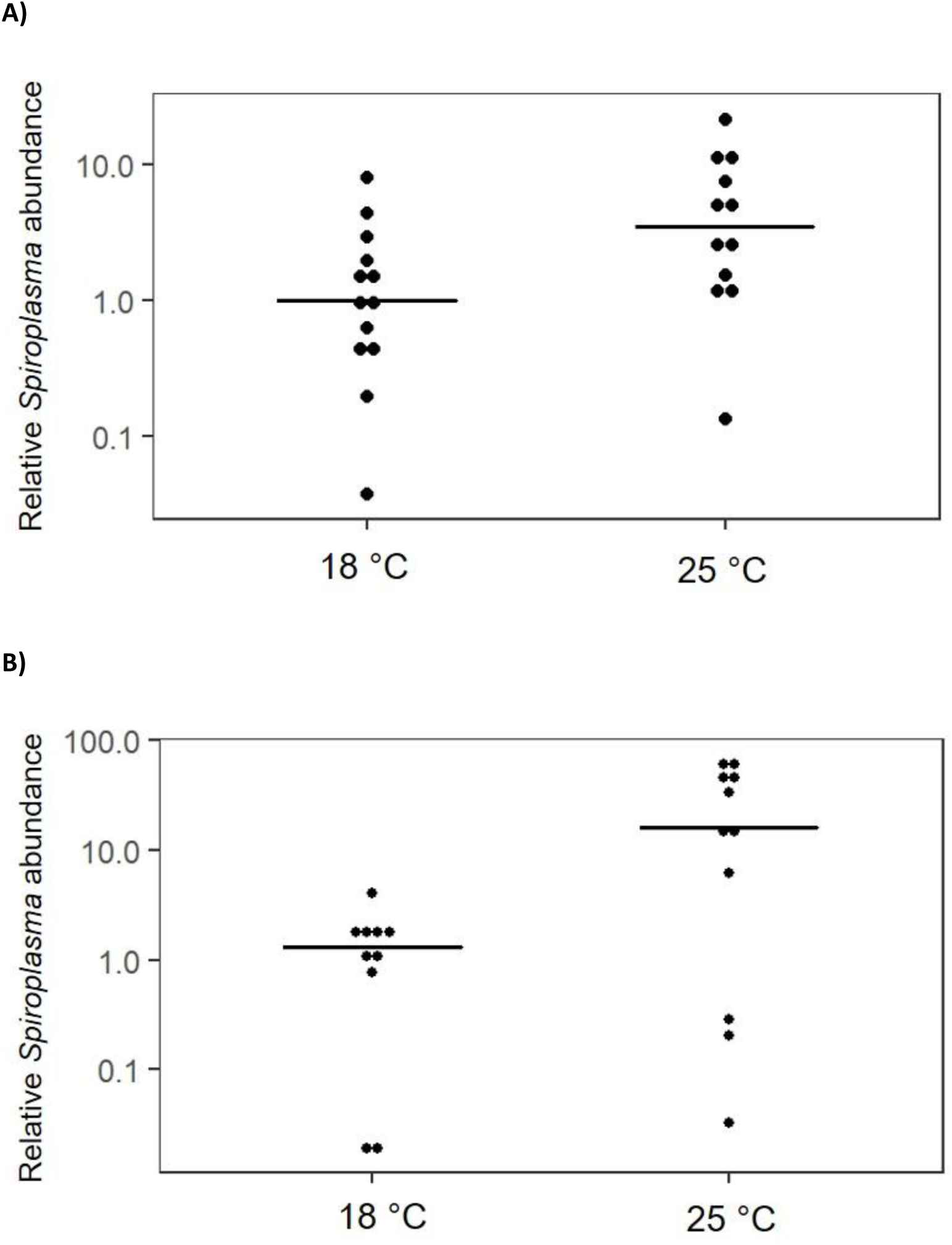
Relative *Spiroplasma* abundance in *D. hydei* pupae reared at 18 or 25 °C, respectively. The two panels represent different blocks in which the experiment was conducted. Each dot represents a single pupa and lines represent medians of the replicates.

## Discussion

The impact of thermal environment on the capacity for a parasite to infect, its virulence and onward transmission are well appreciated within the context of an insect’s standard immune response (Thomas and Blanford, 2003). Recently, it has been understood that heritable microbes – protective symbionts – form an important additional component of defence in many insects (Vorburger and Perlman, 2018). In this study, we investigated the impact of temperature on the capacity of a symbiont-bearing host to complete development following wasp attack, and on the rate at which the symbiont infects progeny through vertical transmission. We examined these parameters in the *D. hydei*-*Spiroplasma* model of protective symbiosis and observed both aspects – protection and vertical transmission rate – were affected by thermal environment. In both cases, *Spiroplasma* performance was impeded in cooler environments, with loss of protection at 18 °C, and increased rates of segregational loss at 15 °C. We conclude this protective symbiosis is strongly thermally sensitive.

Treatments of protective symbiosis in insects have commonly concluded the presence/absence of a protective phenotype from laboratory study under one thermal environment. The present study indicates thermal environment is key in determining the presence/absence or quantitative level of a protection phenotype. We cannot therefore infer the presence or degree of protection in nature from the observation of a symbiont strain that has been previously studied only at one particular laboratory temperature. Further, any account of ‘real world’ dynamics of the tripartite interaction – both in terms of *Spiroplasma* population biology, host-parasite dynamics, and host-parasite coevolution, requires evaluation beyond standard simplified lab conditions. Our data indicate protection would only regularly occur in the very warmest months of the year in the UK, even if the symbiont is present, although this could be extended in composting environments that generate internal warmth. This seasonality of protection is an important consideration not only for symbiont dynamics, but also for the host-parasite ecological dynamic, and for the role of alternate protection methods, such as encoded in immune defences (Vorburger and Perlman, 2018). An expectation is that protection that is environmentally contingent would broaden conditions for the retention of alternate protection systems that are effective during otherwise unprotected periods.

The likely consequence of cool environment-induced instability of the symbiosis is latitudinal, altitudinal and potentially seasonal variation in *Spiroplasma* frequency. Whilst it represents a single datum, it is notable that the Tunbridge Wells population reported in this paper is both the most northerly screened for *Spiroplasma* to date and has low prevalence (15%) compared to other populations. These data recapitulate the *Spiroplasma* – *D. melanogaster* symbiosis. Here, the male-killing phenotype/vertical transmission are even more cool sensitive, with loss of vertical transmission at 18 °C (Montenegro and Klaczko, 2004), and the symbiosis has to date been recorded only in the tropics (Montenegro *et al.*, 2005; Pool *et al.*, 2006).

The potential reasons for loss of protection at lower temperatures in this symbiosis are unclear currently. Our study emulates the symbiosis and attack process over the host lifecycle, and thus multiple features of the interaction differ between temperatures. The parent flies were allowed to develop at the two focal temperatures and produce progeny which are then attacked over their susceptible period at these temperatures, and resistance (or absence) occurs at the same temperature. Previous studies have noted that *Spiroplasma* titre of flies bred at 18 °C for a single generation was one tenth of that at 25 °C, and observations of our focal strain confirms this. It is commonly observed that the penetrance of symbiont phenotypes is related to titre, with both male-killing efficiency and cytoplasmic incompatibility strength declining as symbiont titre declines (Corbin *et al.*, 2017), and symbiont titre may partly underpin the differences in capacity to rescue wasps.

The fraction of flies attacked at 25/18 °C were similar, as evidenced by fly survival below 20% in the absence of the symbiont in both conditions; however, rates of multiparasitism were not measured. There was evidence that endogenous defence against wasps were thermally sensitive: wasps were more likely to complete development in non-symbiont infected flies at 18 °C than 25 °C. Thus, the endogenous capacity of wasps to complete development declines at 25 °C (mirroring findings in other host species: Ris *et al.*, 2004). The higher fly survival at 25 °C than 18 °C may thus be a product of reduced endogenous defences at 25 °C in combination with any additional symbiont-mediated protection associated with higher *Spiroplasma* titres.

In our study, we observed greater tolerance of *Spiroplasma* transmission to cool temperatures than observed in previous study. Past work on the interaction in a Japanese *Spiroplasma*-*D. hydei* combination indicated loss of all vertical transmission following a single generation reared at 15 °C, and significantly reduced vertical transmission at 18 °C (Osaka *et al.*, 2008). Our study notes reduced vertical transmission, occurring over two generations of passage, at 15 °C and no excess segregational loss was observed at either 18 °C or over 18/15 °C diurnal cycles. The impact of this increased robustness to low temperatures will be to stretch the climate envelope in which the symbiosis can persist. Most importantly, it allows persistence over a long breeding season, rather than truncation loss that might be expected in early or late breeding individuals in a temperate climate like the UK.

It is unclear whether the difference in thermal sensitivity found between this study and that of previous studies has a biological (different strains of host and symbiont) or technical source. In both studies, standard PCR was used to assay *Spiroplasma* presence. This was performed on flies harvested as they appeared in the study of Osaka *et al*. (2008), but on flies allowed to return to 25 °C for 14 days to establish higher titre in our study, following the recovery protocol of Montenegro and Klaczko (2004). False negative PCR assay for symbionts are known to occur more commonly at low titre and have been observed to produce false positives for thermal curing in past work on *Wolbachia* (e.g. Van Opijnen and Breeuwer, 1999). Biological differences are also plausible. In our study, the *Spiroplasma* in the strain/fly combination from the UK was more temperature sensitive than that from Mexico.

Finally, one can conjecture whether the reduction in *Spiroplasma* titre and phenotype at low temperatures is due to a physiological constraint that limits this symbiosis or an adaptation to seasonal wasp attack. Reduced titre may be accompanied by reduced metabolic cost of carrying *Spiroplasma*. The reduced cost comes at a loss of parasite protection phenotype – but it is possible that the times of year when parasite attack is common are those times when temperature is high and thus protection is high (and reciprocally, titre is low when temperature is low and parasite attack rare). In this model, shifting titre is not disadvantageous, and may potentially represent an adaptation. It would be interesting to examine whether the time of year of enemy attack and *Spiroplasma* tolerance to cool temperatures were related across species.

## Acknowledgements

We would like to thank Dr Fabrice Vavre for providing the *Leptopilina heterotoma* (Lh-Fr) strain. We thank Krzysztof Kuś (Oxford University) for providing the qPCR analysis script. We acknowledge funding support from the NERC (doctoral training grants to CC, JJ) and a European Union’s Horizon 2020 Research and Innovation Program under Marie Sklodowska-Curie grant agreement 794507 to EC.

## Data accessibility

Raw data underpinning this study, script for conversion of qPCR raw data, and processed qPCR relative titres, are available in figshare (https://doi.org/10.6084/m9.figshare.c.4957382.v1).

## Author contributions

The research was designed by CC and GH with input from AF and EC. Research was performed by CC, JJ and EC. Data analysis was completed by CC, EC and AF. The paper was written by GH, CC with input from all authors.

## Supplementary material

**Table S1:**
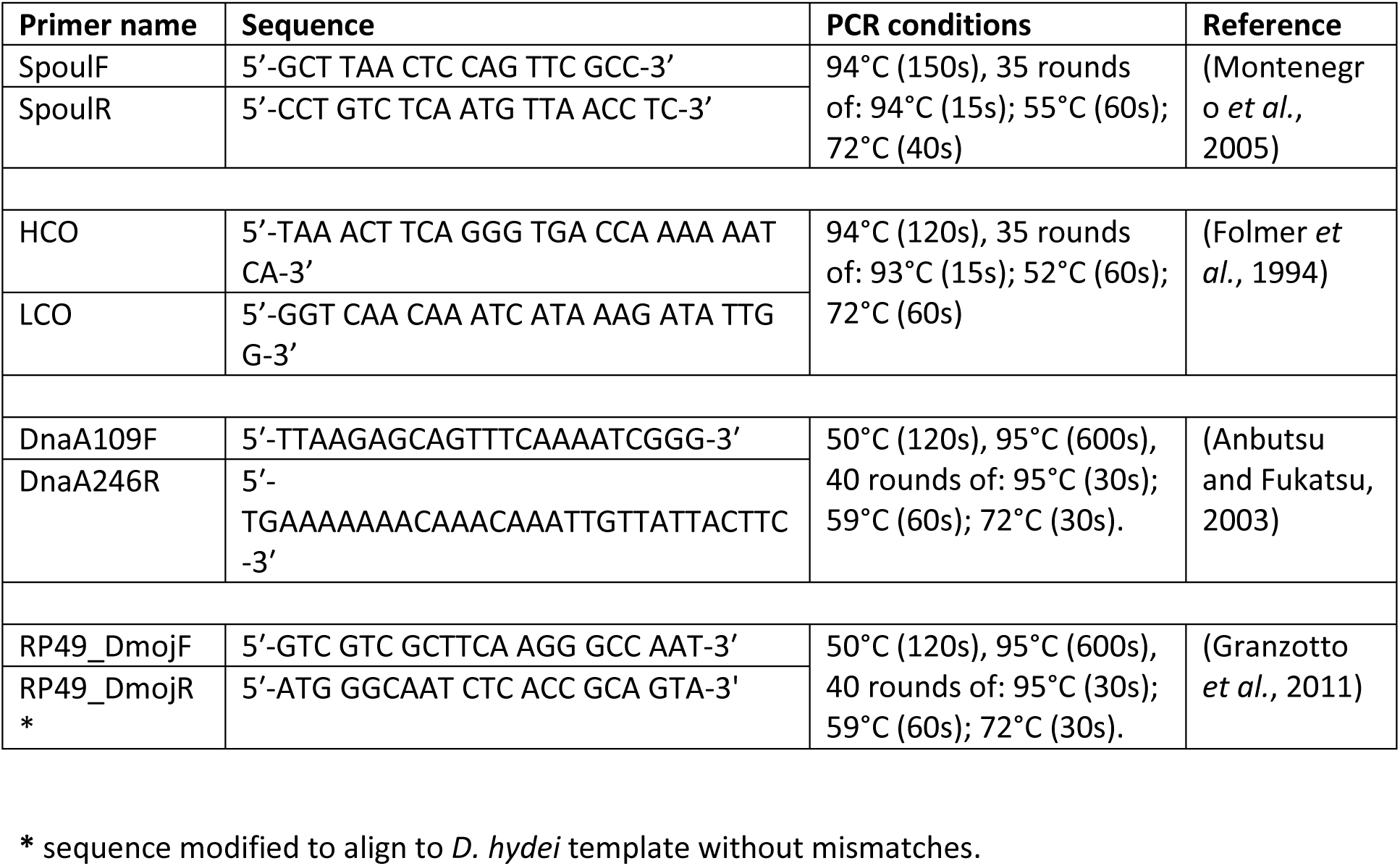
Primer sequences and PCR conditions used in the study.

**Table S2:** GLM results for thermal impacts on *Spiroplasma* transmission.

|  | Estimate | Std. Error | z | Pr(>\|z\|) |
| --- | --- | --- | --- | --- |
| (Intercept) | 11.2213 | 2.8174 | 3.983 | 6.81e-05 |
| Temperature | -2.9336 | 0.7284 | -4.028 | 5.63e-05 |
| Strain | -7.8342 | 2.9268 | -2.677 | 0.007435 |
| Temp x strain | 2.6349 | 0.7912 | 3.330 | 0.000868 |

**Table S3:** GLM results for thermal impact on fly survival in the presence and absence of wasps, and presence/absence of *Spiroplasma* symbiont infection.

|  | Estimate | Std. Error | z | Pr(>\|z\|) |
| --- | --- | --- | --- | --- |
| (Intercept) | 5.39285 | 1.73398 | 3.110 | 0.00187 |
| Temperature | -0.06653 | 0.07566 | 0.879 | 0.379 |
| Spiroplasma presence | -5.88897 | 2.16757 | -2.717 | 0.00659 |
| Wasp attack | -5.97912 | 0.50491 | -11.842 | < 2e-16 |
| Temperature x *Spiroplasma* presence | 0.32129 | 0.09870 | 3.255 | 0.00113 |

**Table S4:**
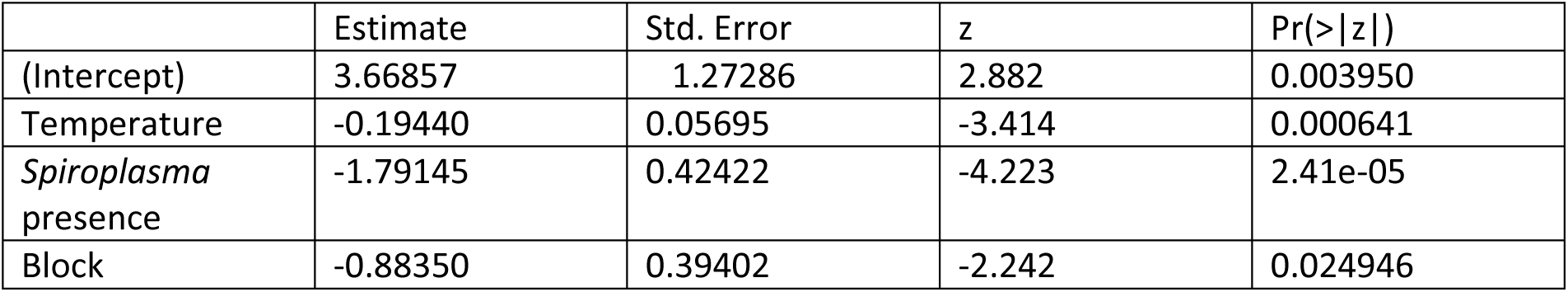
GLM results for thermal impact on wasp survival in the presence and absence of *Spiroplasma* symbiont infection

